# Increased Exertion Variability is Linked to Disruptions in Effort Assessment in Multiple Sclerosis

**DOI:** 10.1101/2025.07.16.665234

**Authors:** Michael H. Dryzer, Bardia Nourbakhsh, Jennifer Keller, Vikram S. Chib

**Affiliations:** Department of Biomedical Engineering, Johns Hopkins School of Medicine, 720 Rutland Avenue, Baltimore, MD, 21205, USA; Department of Neurology, Johns Hopkins School of Medicine, 600 North Wolfe Street, Baltimore, MD, 21287, USA; Kennedy Krieger Institute, 716 North Broadway, Baltimore, MD, 21205, USA; Department of Physical Medicine & Rehabilitation, Johns Hopkins School of Medicine, 720 Rutland Avenue, Baltimore, MD, 21205, USA; Kavli Neuroscience Discovery Institute, 725 N. Wolfe Street, Johns Hopkins University, Baltimore, MD, 21205, USA

**Keywords:** Multiple sclerosis, effort, exertion, motor performance

## Abstract

Accurate assessment of exertion is crucial for determining whether to continue or rest during physical activity. Individuals with multiple sclerosis (MS) often report fatigue and motor impairments, yet the mechanisms underlying their assessments of effortful exertion remain poorly understood. Recent work with healthy individuals suggests that motor variability can distort judgments of effort; however, this relationship has not been explored in individuals with MS. In this study, we investigated how variability in physical exertion affects effort assessment in individuals with MS compared to healthy participants. We had participants exert varying levels of physical effort and retrospectively assess the effort they exerted. Individuals with MS exhibited increased exertion variability and tended to overreport their levels of exertion compared to healthy participants. Increased exertion variability was associated with less accurate effort assessment, and this effect was more pronounced in individuals with more advanced MS. These results suggest a possible mechanism through which motor variability may be associated with inflated perceptions of effort in MS, highlighting a potential account of why efforts feel particularly costly to those individuals living with MS and identifying a promising target for treatment and rehabilitation.

## INTRODUCTION

When performing a physical activity, we must continually assess our effort and use this information to decide whether to persist or take a break. Individuals with multiple sclerosis (MS) consistently report that movement impairments and feelings of physical weakness and fatigue impact their daily lives.^1–4^ Laboratory studies have presented mixed results regarding individuals with MS assessments of effort, showing that after sustained exertion, their perceptions of exertion may be heightened,^5,6^ matched,^7–10^ or even reduced compared to healthy individuals.^11^ These varied findings raise the question of the mechanisms through which physical exertion is translated into judgments of effort in MS.

Several studies have been conducted to investigate the impact of MS on perceived exertion. These studies have involved individuals performing various types of physical exertion (e.g., cycling, arm flexion, finger flexion) and rating their feelings of exertion using self-report scales.^5,7,9,10^ These investigations have reported mixed results regarding the relationships between perceived exertion and MS disease symptomology. One limitation of this previous work has been the use of self-report scales, which provide very coarse subjective measures of perceived exertion. Such scales were designed to quantify the subjective feeling of exertion originating from the sum of all bodily systems, attempting to integrate various components that could lead to feelings of exertion, including signals elicited from peripheral muscles and joints, from central cardiovascular and respiratory functions, and the central nervous system.^12,13^ However, the resulting self-report measures provide only a coarse measure of feelings of effort, making it challenging to assess how different types of effort are perceived objectively and the underlying mechanisms.

Recent work in healthy individuals has begun to identify how the execution of motor performance influences subsequent perceptions of effort. These studies have shown that variability in motor performance is related to judgments of effort, such that increased movement variability reduces an individual’s willingness to perform an action.^14,15^ Works that have directly examined the relationship between physical exertion and subsequent assessments of effort have found that increased exertion variability disrupts participants’ assessments of effort.^16,17^ Across a variety of motor domains, such as gait,^2,18,19^ arm movement,^20–22^ and isometric exertion,^23–25^ individuals with MS exhibit increased motor variability relative to healthy participants. However, the link between motor variability and effort assessments has not been directly explored.

Here, we investigated how variations in motor performance influence assessments of physical effort in patients with MS. We hypothesized that patients with MS would exhibit heightened variability in their physical exertions compared to healthy individuals and that these differences in motor performance would lead to inflated assessments of effort. This hypothesis is informed by studies that have shown increased variability in exertion in MS,^24,26^ as well as evidence indicating that increased exertion variability is associated with higher assessments of effort.^16,17^ Given that deficits in muscle strength and motor performance are associated with MS disease severity,^2,3,24^ we further hypothesized that disruptions in effort assessment in MS would also be related to disease severity. Together, these hypotheses form an account of how pathological declines in motor performance may shape assessments of effort to increase the cost of physical exertion in individuals with MS.

## METHODS

### Experimental setup

The presentation of visual stimuli and acquisition of behavioral data were performed using custom MATLAB (https://www.mathworks.com) scripts implementing the PsychToolBox libraries.^27^ A hand-clench dynamometer (TSD121B–MRI, BIOPAC Systems, Inc., Goleta, CA) recorded grip force exertion. The signals from the sensor were sent to our custom-designed software for real-time visual feedback of participants’ exertions. Participants were instructed to exert a grip force on the sensor in their dominant hand.

### Participants

The Johns Hopkins School of Medicine Institutional Review Board approved this study, and all participants provided informed consent in accordance with the Declaration of Helsinki. Participants were recruited from the Johns Hopkins Multiple Sclerosis Center, and healthy participants were recruited from the Johns Hopkins University community via online postings. Experimenters were not blinded to group participation during data collection. A neurologist verified diagnosis of MS, and a total of 29 persons with MS participated in the study (mean age ± standard deviation, age 46 ± 12y; age range, 23 – 67y; 13 males; median EDSS (IQR), 2.5 (2.5); mean FSS ± standard deviation, 4.92 ± 1.70). Healthy participants were age-matched (± 5y) to the patient cohort and had no history of neurological or neuropsychiatric disorders. Five participants in the control cohort were excluded because they were unable to generate salient associations between physical exertion levels and assessments of effort (*r*-squared between reported and exerted effort during the assessment phase was < 0.5), leaving a final control cohort comprised of 22 participants (mean age ± standard deviation, 46 ± 13y; age range, 23 – 67y; 12 males).

Prior to the experiment, participants were scored on the Expanded Disability Status Scale (EDSS)^28^ and instructed to report the impact of fatigue on their daily life using the Fatigue Severity Scale (FSS).^29^ The EDSS is comprised of a cumulative score reflecting general disability in patients as well as a series of 8 subscores measuring disability in distinct functional systems: pyramidal, cerebellar, brainstem, sensory, bowel and bladder, visual, cerebral (cognitive), and other (reserved for observations that do not fit into any of the other 7 systems) functions. Seven patients completed the cumulative EDSS without reporting functional system subscores, and were excluded from analytical methods that required these subscores, leaving 22 patients for these analyses.

### Experimental paradigm

Before the experiment, participants were informed that they would receive a fixed show-up fee of $30, which was not contingent upon their performance or behavior during the experimental tasks. The effort task used for this paradigm was like those used in previous studies that have examined behavioral and neural representations of effort-based judgements in health and disease.^16,17,30–34^

The experiment began with participants exerting their maximum voluntary contraction (MVC). Participants were unaware of the subsequent experimental phases. They were instructed to squeeze the hand-clench dynamometer with their maximum force over a 4 second duration for each of three consecutive trials, with a 4 second rest period separating each trial. A participant’s MVC was calculated as the maximum force achieved over three consecutive trials.

Participants next underwent an association phase during which they were trained to associate effort levels with the corresponding force exerted on the dynamometer (Fig. 1A). Effort levels were presented on a scale ranging from zero effort units (no exertion) to 100 effort units (80% of a participant’s MVC). Participants proceeded through a randomized order of training blocks, each consisting of five trials at a single target effort level, which varied from 10–80 effort units in increments of 10 effort units. Each association trial began with the numeric presentation of the target effort level (2 s), followed by an effort task with visual feedback in the form of a black vertical bar, similar in design to a thermometer, which increased in white proportional to the level of grip exertion (4 s). The bottom and top of this effort gauge represented effort levels zero and 100, respectively. Participants were instructed to reach the target zone (±5 effort units of the target) as fast as possible and maintain their exertion within the target zone for as long as possible over the course of 4 seconds. Visual indication of the target zone was colored green if the effort produced was within the target zone, and red otherwise. After exertion, if participants remained within the target zone for more than two-thirds of the trial time (2.67□s), the trial was considered a success. Participants received feedback of their success or failure in maintaining the target effort after each trial. To minimize participants’ fatigue, a fixation cross (2–5□s) separated the trials within a training block, and 60□s of rest were provided between training blocks.

**Figure 1.**
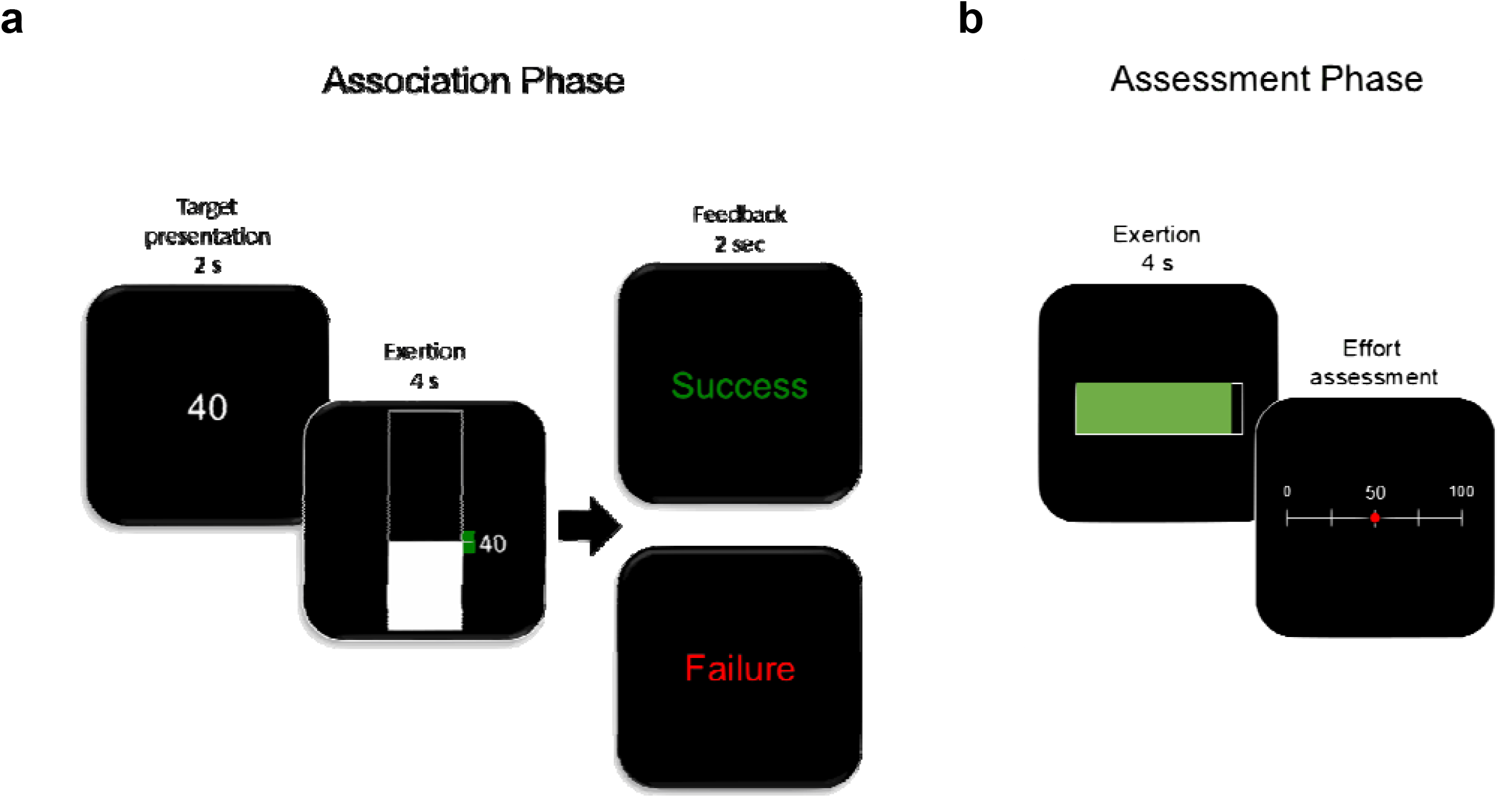
Experimental paradigm. **(a)** During the association phase, participants learned to associate effort levels with force exerted on a hand-clench dynamometer. Effort levels were presented on a scale ranging from 0 (no exertion) to 100 (80% of a participant’s MVC). Each trial began with the numeric presentation of the target effort level, followed by an effort task with visual feedback in the form of a vertical bar that increased in height as force exerted on the dynamometer increased. Simultaneously presented was a target zone that became green when participants reached and maintained the target effort and remained red otherwise. Feedback of task success or failure was presented immediately after completion of each trial. **(b)** During the assessment phase, participants performed an effort assessment task during which they were instructed to exert unknown amounts of effort and assess their levels of exertion. On each trial, participants filled a horizontal bar by exerting on the hand-clench dynamometer. A full bar was representative of an effort level unknown to the participants. The bar turned from red to green once the target effort was reached. Following each exertion, participants estimated their perceived level of exertion using a number line (range 0 to 100). No feedback was provided to participants as to the accuracy of their selection.

Following the association phase, participants performed the assessment phase to evaluate how they generated assessments of their levels of exertion (Fig. 1B). All the effort levels from the association phase (10–80 in increments of 10 effort units) were presented in random order, six times each. Each assessment trial began with the display of a black horizontal bar that participants were instructed to fill by exerting on the dynamometer (4 s). Visual feedback turned red to green once the target effort level was reached. A full bar did not correspond to an effort level of 100 as in the previous phase; here, it represented the target effort level required on each trial. Participants were instructed to reach the target zone as quickly as possible, maintain their force production for as long as possible, and estimate their effort level during exertion. Following the exertion of effort, participants were presented with a number line ranging from zero to 100 and asked to select the effort level they believed they had just exerted. Selection was achieved by moving a computer mouse to the point of selection and clicking the left mouse button to finalize the response. Participants had 4 seconds to make this effort assessment; if they failed to do so, the trial was considered a miss. No feedback was provided to participants regarding the accuracy of their selection. After each selection was made, a fixation cross (2–5□s) appeared on screen to provide a period of rest in between trials. A longer rest period of 60 seconds was provided halfway through the phase.

### Analysis of assessment phase exertions metrics

Assessment phase data was analyzed over the final 1 second of exertion of each trial (referred to as the target period). Data from the first 3 seconds of each trial were excluded to mitigate the effects of force ramping on performance and to account for differences in reaction times. Trials in which participants exerted minimal effort (< 5 effort units) during the target period were excluded from the analysis. We have used a similar approach in previous studies using the same behavioral paradigm.^16,17^

The mean exertion (ME) over the target period of each trial was calculated for all participants. The exertion variability (EV) of each trial was acquired by calculating the standard deviation of exertion during the target period. The normalized exertion variability (EVN) of each trial was calculated by dividing the standard deviation of exertion during the target period by the corresponding mean exertion (i.e., the coefficient of variation of the exertion variability). Normalizing the exertion variability controlled for the correlation between increasing levels of exertion and exertion variability, allowing us to capture the relationship between trial-to-trial variations in exertion variability and assessments of effort. Absolute assessment error (AE) was obtained by calculating the magnitude of the difference between effort assessment (EA) and mean exertion during the target period. Reaction time (RT) was calculated by identifying the timepoint at which exertion surpassed 5 effort units.

### Hierarchical linear modeling of exertion-dependent variations in effort assessments

Hierarchical linear models were used to evaluate the relationships between trial-to-trial variance in exertion performance and assessments of effort, and to establish how this relationship changed in patients with MS. This was accomplished using the *lme4* R package.^35^ We tested relationships between EV, EA, and AE. To compare participant cohorts, we designed hierarchical models with the population tag (control cohort versus MS cohort), appropriate covariate (ME, EV, EVN), and the interactions between these terms as fixed effects. For the random effects, we included participant as a random intercept and the relevant exertion metric of each model as a random slope. All analyses were performed using only assessment phase data, and regressors were z-scored before being input into the models.

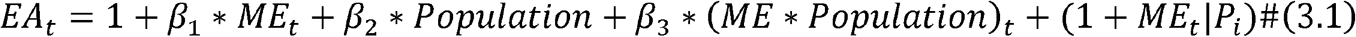

To model the relationship between a participant’s exertions and their effort assessments, we computed the above model where *t* denotes the trial number, *i* denotes the subject number (1– 51), *Population* is a categorical regressor (0 – Control, 1 – MS), and *p*_*i*_ is a categorical identifier for each participant.

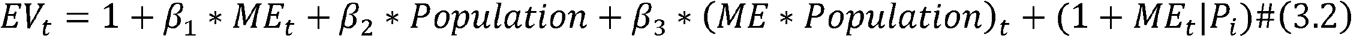

The relationship between a participant’s mean exertion and exertion variability was modeled using the above equation.

Additionally, we designed a model estimating the relationship between a participant’s AE and EVN. This allowed us to test if performance variability influenced errors in assessments of effort.

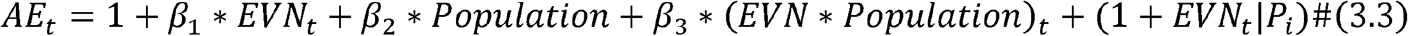

Finally, we conducted a series of models testing for between-patient differences in the relationship between exertion variability and assessment error. To accomplish this, we replaced the categorical *Population* regressor in equation 3.3 with the continuous *EDSS* regressor representing the degree of disability due to MS, yielding the equation presented below. This regressor accepts either the cumulative EDSS score or any one of the functional system subscores, thus, it represents the influence of general disability or disability in a specific functional system. We chose the cerebellar, pyramidal, and sensory functional system scores for our analyses, because each of these systems were relevant in the execution and assessment of physically effortful exertions.

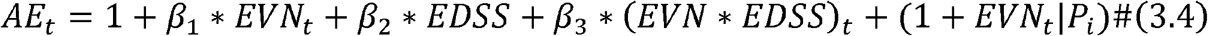

We also adapted the model above to estimate the impact of fatigue severity on assessments of effort, replacing the *EDSS* regressor with *FSS*, which signifies the severity of fatigue.

## RESULTS

Both healthy individuals and those with MS were able to exert and maintain effort at the target levels during the assessment phase (Fig. 2A, B), and there were no differences in MVC between groups (average difference in MVC: 25.05 N; two-sample, two-tailed t-test, *t* = −0.78, *df* = 49, *p* = 0.44). A trial-by-trial analysis of the relationship between exertion variability and mean exertion showed that the variability increased with increasing exertion for both control and MS participants (Fig. 2C; hierarchical linear model, *β* = 0.32, 95% *CI* = [0.08, 0.56], *t* = 2.64, *df* = 2,316, *p*< 0.05). This result is consistent with previous experiments studying isometric force production, which demonstrated that variability in motor output increases in proportion to the amount of exertion (i.e., signal-dependent noise).^36–38^ Comparing the relationship of mean exertion and exertion variability between the control and MS group, we found that participants with MS exhibited a greater increase in exertion variability as exertion increased (hierarchical linear model, *β* = 1.00, 95% *CI* = [0.32, 1.68], *t* = 2.87, *df* = 2,316, *p* < 0.01). These results align with our hypothesis of MS being associated with increased exertion variability during effortful motor output.

**Figure 2.**
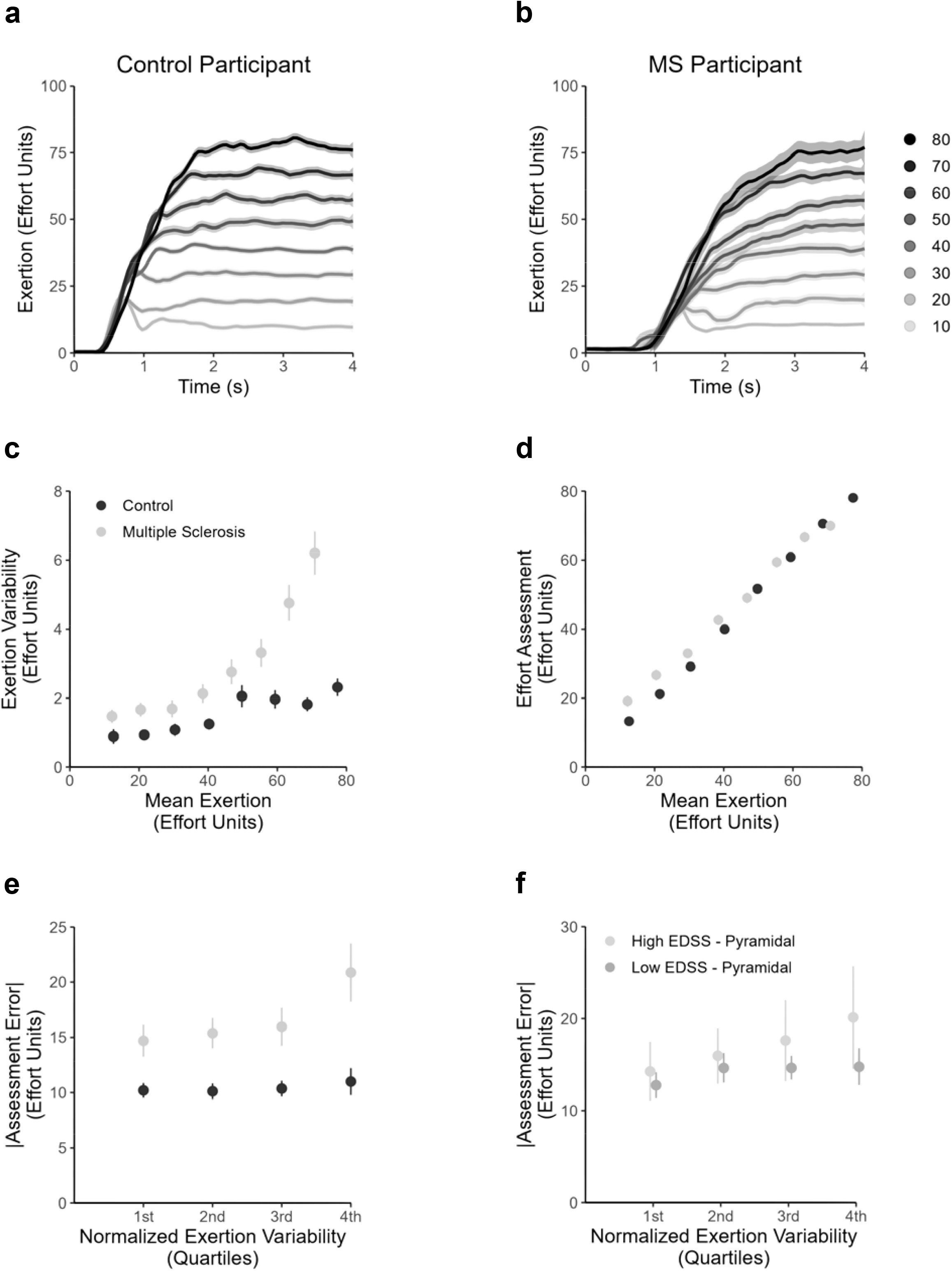
Behavioral results (n = 51) **(a, b)** Assessment phase mean exertion trajectories from representative participants in the control (left) and MS (right) cohorts. Data are presented in effort units, which were relative to a participant’s MVC, and were averaged within each target effort level. Both groups of participants were able to reach and maintain exertions at the target effort level. Patients with MS had significantly higher reaction times to trial onset compared to controls (average difference in reaction time: 0.40 s; two-sample, two-tailed t-test, p « 0.001). The remaining plots (**c** – **f**) were used for illustration purposes and not statistical inference, which was accomplished with hierarchical linear models in R. All calculations used data in the target period (final 1 s of each trial). (Control group – black circles; MS group – gray circles.) **(c)** Exertion variability as a function of mean exertion. Individuals with MS had a more pronounced relationship between increases exertion variability with increasing levels of effortful exertion (hierarchical linear model, *β* = 1.00, 95% *CI* = [0.32, 1.68], *t* = 2.87, *df* = 2,316, *p* < 0.01). Error bars are SEM. **(d)** Effort assessment as a function of mean exertion. Individuals with MS overestimated levels of exertion relative to healthy individuals (hierarchical linear model, *β* = 2.65, 95% *CI* = [4.17, 1.13], *t* = 3.40, *df* = 2,316, *p* < 0.01). Error bars are SEM. **(e)** The magnitude of assessment error as a function of normalized exertion variability. Assessment errors were acquired by calculating the difference between assessed effort and the mean exertion on each trial. Normalized exertion variability, which accounts for performance-related increases in exertion variability, was calculated by dividing exertion variability by mean exertion for each trial. For illustration, assessment errors and normalized exertion variability were pooled in quartiles of normalized exertion variability. Individuals with MS exhibited more pronounced increases in assessment error with increases in exertion variability (hierarchical linear model, *β* = 3.16, 95% *CI* = 1.34, 4.98, *t* = 3.40, *df* = 2,316, *p* < 0.01). Error bars are SEM. **(f)** Individuals with greater pyramidal MS-related disability exhibit increased assessment error. For illustration, assessment errors and normalized exertion variability were pooled in quartiles of normalized exertion variability. Individuals with high Pyramidal Expanded Disability Status Scale (EDSS) scores exhibited a more pronounced relationship between normalized exertion variability and errors in effort assessment (hierarchical linear model one-tailed t-test, *β* = 1.49, 95% *CI* = 0.16, ∞, *t* = 1.82, *df* = 1,266, *p* < 0.05). Errsor bars are SEM.

Across groups, we found that assessments of effort increased with mean exertion (Fig. 2D; hierarchical linear model, *β* = 18.89, 95% *CI* = [17.37, 20.40], *t* = 24.37, *df* = 2,316, *p* « 0.001), confirming that participants had a good understanding of the task. We found a significant interaction between mean exertion and group, indicating that individuals with MS exhibited increased overall assessments of effort compared to control participants (hierarchical linear model, *β* = −2.65, 95% *CI* = [−4.17, −1.13], *t* = −3.40, *df* = 2,316, *p* < 0.01), and this interaction was predominantly driven by over assessment of effort at lower exertion levels, and under assessment at higher exertion levels.

Next, we tested for an effect of mean exertion on assessments of effort, reasoning that increased exertion variability may distort the perception of effort production and influence assessments of effort. To test our hypothesis regarding the influence of MS on the relationship between exertion variability and assessments of effort, we compared metrics related to exertion performance and participants’ ratings of exertion. For each recall trial, we calculated a measure of normalized exertion variability as the standard deviation of exertion divided by the mean exertion on that trial (i.e., the coefficient of variation). This normalized metric allows us to account for the expected increase in variability with exerted effort (Fig. 2C) and evaluate how trial-to-trial variations in exertion variability are related to effort assessment errors (absolute difference between measured mean exertion and reported effort).^16,17^

Across groups, participants’ normalized exertion variability predicted the magnitude of errors in effort assessment (Fig. 2E; hierarchical linear model, *β* = 2.38, 95% *CI* = [1.20, 3.57], *t* = 3.94, *df* = 2,316, *p* < 0.001). This indicated that both groups of participants (control: one-tailed hierarchical linear model, *β* = 1.43, 95% *CI* = [0.18, ∞], *t* = 1.89, *df* = 1,048, *p* < 0.05; MS: hierarchical linear model, *β* = 3.16, 95% *CI* = [1.39, 4.95], *t* = 3.54, *df* = 1,268, *p* < 0.01) were less accurate in assessing their levels of exertion following trials in which they exhibited higher degrees of variability. This finding aligns with previous studies which have shown that increased exertion variability inflates assessments of effort.^16,17^ Furthermore, we found that this relationship was more pronounced in individuals with MS (hierarchical linear model, *β* = 3.16, 95% *CI* = [1.34, 4.98], *t* = 3.40, *df* = 2,316, *p* < 0.01), suggesting that MS influences the degree to which exertion variability impacts the transformation of exertion performance into assessments of effort.

We next evaluated how the relationship between exertion variability and effort assessments modulated individual differences in the degree of MS symptoms. Specifically, we focused on cumulative EDSS scores and the EDSS subscores for the pyramidal, cerebellar, and sensory functional systems. We chose these functional systems because of their importance in executing motor commands and processing of afferent sensory feedback, which are likely critical for effort exertion and perception. Incorporating these scores into our models allowed us to estimate how general disability and disability in specific functional systems mediated the relationship between exertion variability and assessment error.

EDSS scores served as a significant modulator of the relationship between exertion variability and errors in effort assessment, with those individuals with higher EDSS exhibiting a relationship between exertion variability and assessment error (Fig. 2F; hierarchical linear model one-tailed t-test, *β* = 1.49, 95% *CI* = [0.16, ∞], *t* = 1.82, *df* = 1,266, *p*< 0.05). This aligns with our prediction that patients experiencing a higher degree of MS symptomology are more sensitive to variability in motor performance, impacting disruptions in their effort assessments.

When testing for the influence of the pyramidal (hierarchical linear model, *β* = 1.66, 95% *CI* = [0.18, 3.15], *t* = 2.19, *df* = 1,004, *p*= 0.041), cerebellar (hierarchical linear model, *β* = 1.56, 95% *CI* = [0.05, 3.09], *t* = 2.01, *df* = 1,004, *p*= 0.058), and sensory functional systems on the link between assessment error and exertion variability (hierarchical linear model, *β* = 0.98, 95% *CI* = [−0.62, 2.60], *t* = 1.19, *df* = 1,004, *p* = 0.25), we found that only disruptions to the pyramidal functional system predicted increased assessment error through a significant positive interaction with normalized exertion variability. This finding further suggests that increased effort assessment error in MS may be most related to the degree of disability in the pyramidal functional system, which is most related to muscle weakness and limb mobility.

## DISCUSSION

Here, we demonstrate that individuals with MS exhibit increased variability in physical exertion and subsequent assessments of effort, compared to healthy individuals, and that this motor variability contributes to errors in assessments of effort. We show that exertion variability has a greater influence on effort assessment error in individuals who present with more MS disability, especially those experiencing deficits in the pyramidal functional system. Our findings align with previous work, which has shown that increased variability in physical exertion is associated with subsequent inflated assessments of effort.^16,17^ Additionally, our finding of heightened exertion variability aligns with studies of MS that have found patients display higher motor variability than healthy individuals (Crenshaw et al., 2006; Sosnoff et al., 2012; Societ et al., 2013; Carpinella et al., 2014; Pellegrino et al., 2015; Wolkorte et al., 2015; Arpin et al., 2016; Pellegrino et al., 2018).^2,18–24^ Our findings extend beyond these previous works by revealing a mechanism through which exertion variability translates into subjective judgments of effort, and how the interplay between effort exertion and assessment is mediated by MS symptomatology.

The nervous system generates and senses effortful physical exertion through central and peripheral mechanisms. Afferent fibers transmit sensory feedback from muscles, joints, and tendons, which detect muscle actuation and metabolic byproducts related to exertion.^39,40^ This peripheral information is relayed to brain regions, including the somatosensory cortex, insula, and anterior cingulate, contributing to the perception of effort and bodily state.^30,41–44^ Central motor commands originating from the motor cortex contribute to corollary discharge, creating internal signals that inform the brain of intended movement and the expected level of exertion.^40,45–47^ Investigations of motor and sensory deficits in MS suggest that the processes integral to generating corollary discharges and motor commands are affected. The demyelination associated with MS has been shown to compromise the corticospinal tract, which transmits central motor commands to muscles and drives motor output.^24,48,49^ In this context, our findings suggest that participants with more deficits in the pyramidal functional system exhibited greater assessment error, which may be due to the interaction of these deficits with exertion variability.

MS-induced alterations in cognitive and perceptual processes may further influence the relationship between exertion variability and errors in effort assessment. Studies exploring the neural basis of MS have documented weakened intra-cortical connectivity between sensory and motor cortices.^50,51^ This reduction in responsiveness may not only contribute to the higher motor variability observed in individuals with MS but also impair their ability to assess exertion accurately. Concurrently, the pathological fatigue often reported in MS patients correlates with deteriorated interoception, affecting their capacity to monitor bodily sensations and manage exertion effectively.^52,53^

Interoceptive sense may play a crucial role in our findings, as it serves as a framework through which individuals perceive their internal bodily states, including the sensations associated with exertion.^40,41,54^ The impaired interoceptive capacity in individuals with MS may limit their ability to effectively gauge the physical resources required for tasks, further disrupting the relationship between motor variability and effort perception.^52,53,55^ With higher exertion variability, individuals may struggle to accurately integrate sensory information from both their exertion and their internal physiological state, leading to distorted assessments of effort. These impaired perceptions may cause a disconnect between actual exertion levels and perceived effort, contributing to the inflated self-reports of effort we observed. Understanding interoceptive deficits in MS could provide insights into the mechanisms underlying poor effort assessments, highlighting the importance of interoceptive training in conjunction with rehabilitation as a potential avenue for enhancing effort assessment.

Our findings have significant implications for rehabilitation approaches. They underscore the need for clinical assessments to integrate both motor performance metrics and subjective assessments of effort. Recognizing how variability affects effort perception can guide the development of targeted interventions that promote accurate effort estimations, supporting greater engagement in physical activities. Strategies that enhance the consistency of motor output or focus on sensory feedback pathways could be particularly beneficial for managing the effects of heightened motor variability. Moreover, incorporating interventions aimed at improving interoceptive awareness may facilitate better integration of bodily signals, helping individuals form more accurate perceptions of their effort expenditure during exertion.

Our study sheds light on how MS affects individuals’ ability to assess their physical exertion accurately. We found that the extent of MS symptomology significantly modulates the degree to which exertion variability influences judgments of effort. By illustrating a novel mechanism through which variability in motor performance distorts estimations of physical effort, this work adds to the understanding of the interplay between motor control and judgements of exertion in MS.

## ACKNOWLEDGEMENTS

This work was supported by the Eunice Kennedy Shriver National Institute of Child Health & Human Development of the National Institutes of Health under Award Number R01HD097619 and the National Institutes of Mental Health R01MH119086. The authors declare no conflicts of interest.

## Notes

### Competing Interest Statement

The authors have declared no competing interest.

